# Molecular signatures of the rediae, cercariae and adult worm stages in the complex life cycles of parasitic flatworms (Psilostomatidae, Trematoda)

**DOI:** 10.1101/580225

**Authors:** Maksim A. Nesterenko, Viktor V. Starunov, Sergei V. Shchenkov, Anna R. Maslova, Sofia A. Denisova, Andrey I. Granovich, Andrey A. Dobrovolskij, Konstantin V. Khalturin

**Affiliations:** Department of Invertebrate Zoology, St-Petersburg State University. St-Petersburg 199034, Russia; Zoological Institute Rus. Acad. Sci., St-Petersburg 199034, Russia; The A.O.Kovalevsky Institute of Marine Biological Research of RAS, Sevastopol 299011, Russia; Marine Genomics Unit, OIST, 1919-1 Tancha, Onna-son, Kunigami-gun, Okinawa, 904-0495 Japan

**Author notes:** Email addresses: MAN, VVS, SVS, ARM, SAD, AIG, AAD, KVK.

## Abstract

Trematodes are one of the most remarkable animals with complex life cycles with several generations. Life histories of a parasitic flatworms include several stages with disparate morphological and physiological characteristics follow each other and infect hosts ranging from mollusks to higher vertebrates. How does one genome regulate the development of various life forms and how many genes are needed to the functioning of each stages? How similar are molecular signatures of life stages in closely related species of parasitic flatworms? Here we present the comparative analysis of transcriptomic signatures of the rediae, cercaria and adult worm stages in two representatives of the family Psilostomatidae (Echinostomata, Trematoda) - *Psilotrema simillimum* and *Sphaeridiotrema pseudoglobulus*. Our results indicate that the transitions between the stages of the complex life cycle are associated with massive changes in gene expression with thousands of genes being stage-specific. In terms of expression dynamics, the adult worm is the most similar stage between *Psilotrema* and *Spaeridiotrema*, while expression patterns of genes in the rediae and cercariae stages are much more different. This study provides transcriptomic evidences not only for similarities and differences between life stages of two related species, but also for cryptic species in *Sphaeridiotrema*.

## Introduction

Life cycles with morphologically distinct stages occur in various clades of the animal kingdom ^1^. They are typical for free-living Cnidaria ^2^ and Aphiodioidea ^3^ as well as for parasitic Apicomplexa ^4^ and Trematoda ^5^. In contrast to simple life cycle that include only one ontogeny, the complex life cycles are characterized by alteration of several generations and each of them has its own ontogeny ^6^.

Trematoda is a clade of parasitic flatworms that possess one of the most striking examples of complex life cycles ^7^. In the most common variant, the trematode life cycle include three hosts: an invertebrate animals (usually a gastropod mollusk) and a vertebrate as intermediate and definitive hosts, respectively ^5^. Adult worm that inhabits definitive host is usually hermaphroditic and lays eggs from which a free-living larva of the next parthenogenetic generation, called the miracidium, hatches. The main goal of miracidia is to find and infect the first intermediate host. Inside of the intermediate host it transforms into an individual of the next generation. In “redioid” species it turns into a rediae, and in “sporocystoid” species it gives rise to the sporocysts which grow through the tissues of a host. After several cycles of self-reproduction, the individuals of the parthenogenetic generation produce cercariae – the free-living larvae of the amphimictic generation. Cercaria leave the first intermediate host and spread around in search for a new host. If cercariae manages to infect the right definitive host it transforms into an adult worm that again produces eggs with future miracidia, thereby completing the life cycle.

Trematodes of medical or veterinary importance include *Paragonimus westermani* (human lung fluke), *Clonorchis sinensis* (human liver fluke), *Fasciola hepatica* (cattle liver fluke) and, most importantly, the schistosomes (blood flukes) ^8^. *Schistosoma japonicum* ^9^ and *Schistosoma mansoni* ^10^ were the first representatives of trematodes with the sequenced genomes. Availability of genomic data allowed to study many aspects of Trematode biology at the molecular level, including differential gene regulation and epigenetics ^11^. Nevertheless, the blood flukes possess numerous traits that make them very different from the rest of Trematoda, for instance the presence of a schistosomula stage and two separate sexes in their life cycle suggest their highly specialized and evolutionary derived state.

The development of the New Generation Sequencing (NGS) has led to increase of transcriptomic data from different trematode species (^12,13,22,23,14–21^). However, the majority of studies describes only one stage of the life cycle, mostly adult worms (^12–20^). Only few studies provide the comparative analysis of the different stages of the same generation (^22,23^) or the same stage in different species (^19–21^). Comparative transcriptomic analysis of different generations within one life cycle has not been performed so far in any species other than schistosomatides. Such experiment are impeded by a number of the technical challenges. It is prohibitively difficult to maintain complex life cycles in laboratory conditions as that requires cultivation of multiple intermediate and definitive hosts. However, comparative data of that kind are critically important for the understanding of the origin and evolution of trematode life cycles and could provide new insights for treatment and prevention of human and animal trematodosis.

The representatives of the family Psilostomatidae have a dixenous life cycle which appear to be the closest to the “archetypical” life cycle according to modern ideas about the digeneans evolution ^5^. The free-swimming cercariae of these parasites are encysted outside the body of their intermediate host, but in close connection with it (on the mollusk shells or under their mantle fold). The adult psilostomatid worms are typical histiophages and hermaphrodites that parasitize in the intestines of birds. The adult worm body does not have any secondary modifications and possesses a shortened uterus. This feature is useful for experimental work since it allows to reduce the level of adult worm tissue contamination by developing miracidiae. Free-swimming miracidiae actively infect the first intermediate host and give rise to a typical microhemipopulation of rediae. The rediae have a well-developed gut and an extensive brood cavity with developing cercariae. Psilostomatidae use the same species of prosobranch snails, *Bithynia tentaculata*, as the first and the second intermediate hosts, which facilitates their maintenance in the laboratory.

In this paper, we present the comparative transcriptomic analysis of the rediae, cercariae and adult worms of the two species of Trematoda, belonging to the Psilostomatidae family – *Psilotrema simillimum* (Mühling, 1898) and *Sphaeridiotrema pseudoglobulus* (McLaughlin, Scott, Huffman, 1993). The aim of our research was to identify genes with stage-associated expression which most probably contribute to disparate anatomical and physiological characteristics of the life stages. Another goal was to determine similarities and differences in gene expression patterns within one life cycle and between two related species of trematodes.

## Results

### Animal cultivation and library preparation

*Sphaeridiotrema pseudoglobulus* (McLaughlin, Scott, Huffman, 1993) and *Psilotrema simillimum* (Mühling, 1898) have dixenous life cycles (only two hosts). The mollusk *Bithynia tentaculata* (Linnaeus, 1758) is an intermediate host, and waterfowl birds represent the definitive hosts (Fig.1). Naturally infected mollusks were collected from water parts of plants and stones of the Kristatelka pond (Peterhof, Russia) and maintained at the Dept. of Invertebrate Zoology SPbSU. The identification of the cercariae species was carried out using a light microscope according to the morphological features. Rediae and cercaria stages were obtained from cultivated mollusks and the adult worms were cultivated in the experimentally infected chicken.

For both *Psilotrema simillimum* and *Sphaeridiotrema pseudoglobulus* at least 23.3 million pairs of reads were obtained for each of the two biological replicates of the redia, cercaria, and adult worm stages (Tab.1). Contamination with host-derived sequences did not exceed 6.3% from the total number of reads in each library. More than 70% of reads remained in all the libraries after the removal of potential host-derived sequences, adapters and poor-quality reads.

### Quality and completeness of de novo assembled transcriptomes

From BUSCO database representing single-copy orthologs of Metazoa 89.4% (Complete: 83.6%; Fragmented: 5.8%) and 90.7% (Complete: 86.6%; Fragmented: 4.1%) are present in *P.simillimum* and *S.pseudoglobulus* reference transcriptomes, respectively (Tab.2). Among the 79 orthologs missing in our assemblies, 67 sequences were also not found in the previously published genomes and transcriptomes of Trematoda. Thus, most probably these genes represent lineage-specific gene losses most probably associated with the parasitic life style. The complete list of the “common” missing genes with their annotations are presented in Table S2.

TransRate score of assemblies were equal to 0.237 for *P.simillimum* and 0.2095 for *S.pseudoglobulus*. 117130 and 146563 contigs with highest scores were used in the downstream analysis, which comprise 72% of the total number of assembled sequences from *P.simillimum* and *S.pseudoglobulus*, respectively.

### Annotation and orthologues searching

According to ESTScan prediction, 21075 (18%) and 49626 (34%) of transcripts in the assemblies of *P.simillimum* and *S.pseudoglobulus* had open reading frames (ORFs) with more than 70 amino acids, respectively. 62% of predicted protein in both species match to the sequences from Digenea or did not have any matches in NR NCBI protein database. Latter category most probably represents the novel sequences of Trematoda that are strongly derived in Psilostomatidae.

Among assembled transcripts of *P.simillimum* and *S.pseudoglobulus* 17225 and 39346 sequences, respectively, have expression levels above 1 transcript per million (TPM), which constitute 85% and 74% of the reference transcripts of the corresponding species. In both species, the open reading frame was found in approximately 70% of the sequences with expression levels equal to or greater than 1TPM. Gene Ontology annotations could be assigned to 9947 and 22554 proteins in *P.simillimum* and *S.pseudoglobulus*, respectively (Table S12-13). KOBAS 3.0 results showed that 8125 proteins of *P.simillimum* had matches in the metabolic pathways of *H.sapiens* and 9978 had matches in *S.mansoni*. In *S.pseudoglobulus*, 17769 proteins could be assigned to human pathways and 22763 proteins had clear counterparts among the proteins of *S.mansoni*.

### Gene sets in Plathelminthes

Comparison with *Schmidtea mediterranea, Opisthorchis viverrini, O.felineus, Clonorchis sinensis, Fasciola hepatica, Schistosoma mansoni, Trichobilharzia regenti* proteomes allowed to identify 11138 groups of orthologous proteins (Table S3). As shown in Fig.1H, three clusters of species are obvious based on the orthologs number shared among them. One cluster contains three species of Opistorchidae: *O.felineus, C. sinensis* and *O. viverrini*; second one unites schistosomatides: *S. mansoni* and *T.regenti. Fasciola hepatica* groups together with *P.simillimum* and *S.pseudoglobulus*. Only 3075 groups of orthologous proteins (OGs) include sequences from all nine species analyzed and 121 OGs are species-specific (Fig. 1H, Table S3). Complete statistics for all the species are presented in Table S4 (see Supplementary materials).

**Figure 1:**
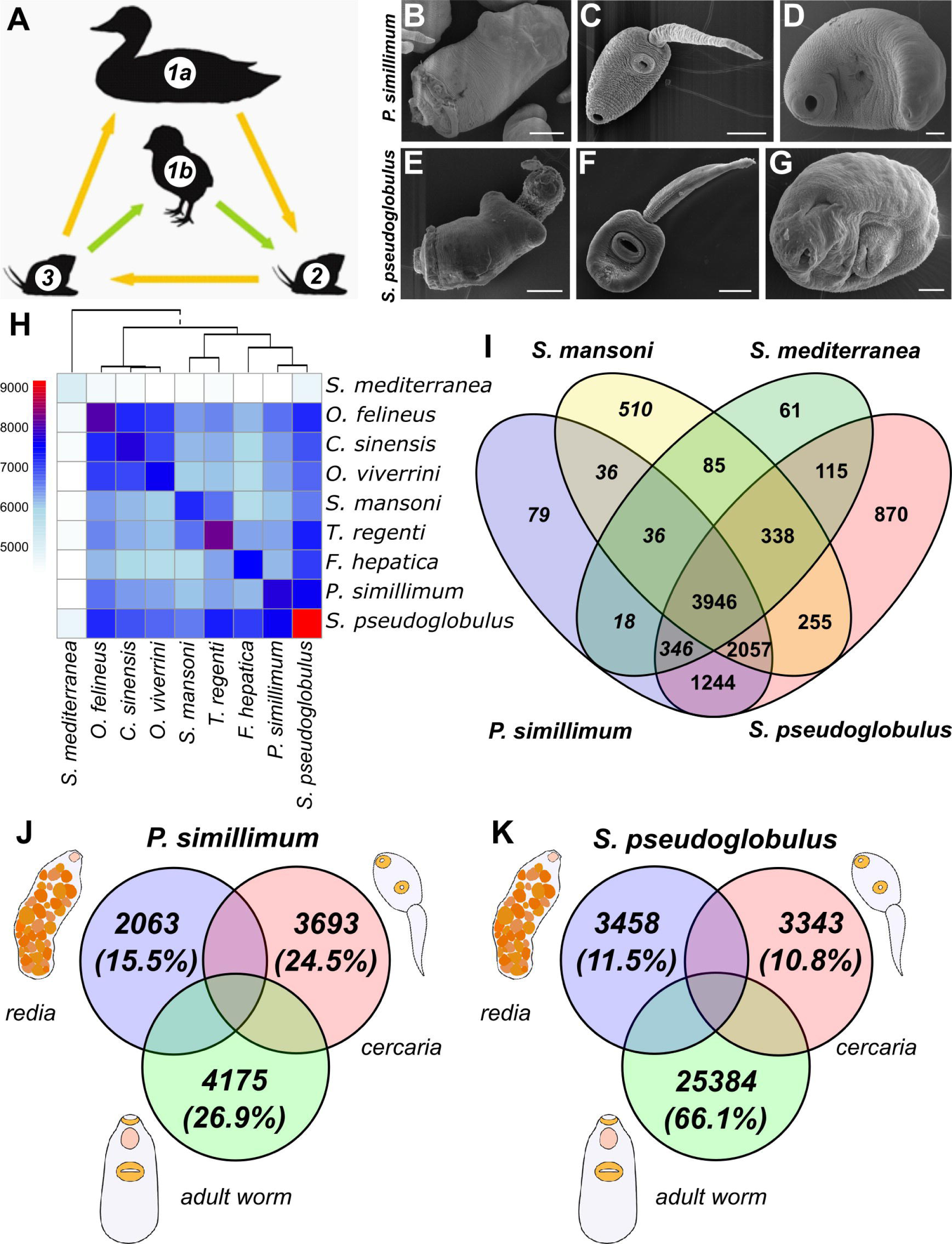
Complex life cycle, similarities and differences between life cycle stages and Trematodes species. **(A)** The realization of Psilostomatidae complex life cycle in wild (yellow arrows) and laboratory (green arrows) conditions; 1a – Definitive host in the wild (waterfowl birds), 1b – Definitive host in the lab (chicken), 2 and 3 – first and second intermediate hosts (mollusk, *Bithynia tentaculata*). **(B-G)** Scanning electron microscope of *Psilotrema simillimum* **(B-D)** and *Sphaeridiotrema pseudoglobulus* **(E-G)** life cycle stages; **(B, E)** – rediae (parasitic stage of parthenogenetic generation), **(C, F)** – cercariae (free-living larvae of amphimictic generation), **(D, G)** – adult worms of amphimictic generation. **(H)** Number of shared orthogroups between *Schmidtea mediterranea, Opisthorchis felineus, O.viverrini, Clonorchis sinensis, Schistosoma mansoni, Trichobilharzia regenti, Fasciola hepatica, Psilotrema simillimum* and *Sphaeridiotrema pseudoglobulus*. Based of gene sets there are three clusters corresponding to the Opisthorchiidae, Schistosomatidae and Echinostomata. **(I)** Venn diagram shows relation between sets of orthogroups, including *Schistosoma mansoni, Schmidtea mediterranea, Psilotrema simillimum* and *Sphaeridiotrema pseudoglobulus*. **(J-K)** Venn diagram shows the number of genes with stage-specific expression in *P.simillimum* **(J)** and *S.pseudoglobulus* **(K).**

The comparison of the orthologous groups (OGs) among four species - *S.mediterranea, S.mansoni, P.simillimum* И *S.pseudoblogulus* revealed that 3946 OGs are common for all studied flatworm species. 2057 OGs are specific for trematodes, and 1244 OGs are specific to the family Psilostomatidae (Fig. 1I). *P.simillimum* has 79 species-specific OGs and in 870 OGs are specific for *S.pseudoglobulus. S.mansoni* also has a lot of species-specific proteins (510 OGs), but their number in *S.pseudoglobulus* clearly stands out. It was an unexpected observation that the number of species-specific OGs in *P.simillimum* or *S.pseudoglobulus* differs more than 9-fold (Fig. 1E).

### Genes with stage-specific expression in rediae, cercariae and adult worms

Each stage of the trematode life cycle is characterized by structures and functions that are not present in the other stages. Thus, different sets of genes must be activated in order to allow these differences. How many genes in the genomes of the trematodes are utilized for generation of stages-specific trait? In order to answer this question, we identified genes with preferential expression in the redia, cercaria and adult worm stages in *Psilotrema* and *Sphaeridiotrema*.

According to the Jongeneels criterion (see Materials and Methods), approximately 15.5% of genes in *P.simillimum* have rediae-specific expression, 24.5% are specific for the cercariae, and 26.9% - for the adult worm. Differences between *S.pseudoglobulus* life cycle stages in terms of gene sets utilized are more pronounced: 10.8% of genes have rediae-specific expression, 11.5% are rediae-specific, and 66.1% of the genes are active mainly on the adult worm stage. In both species examined the majority of genes with stage-specific expression are associated with the adult worms (fig.1J-K) and this tendency is most prominent in *Spaeridiotrema*.

Comparison of gene sets with stage-specific expression revealed that in contrast to *P. simillimum*, where stage-specific sets are most similar to themselves only (Fig.2A), in *S. pseudoglobulus* there is a significant overlap between the genes specifically expressed in rediae and cercariae stages (Fig. 2B). This observation might be explained by the fact that inside of the redia the next generation (cercaria) is developing. This can be clearly observed in Fig.2E where numerous round-shaped structures visible inside of the *Psilotrema* redia are the developing cercaria larvae.

**Figure 2:**
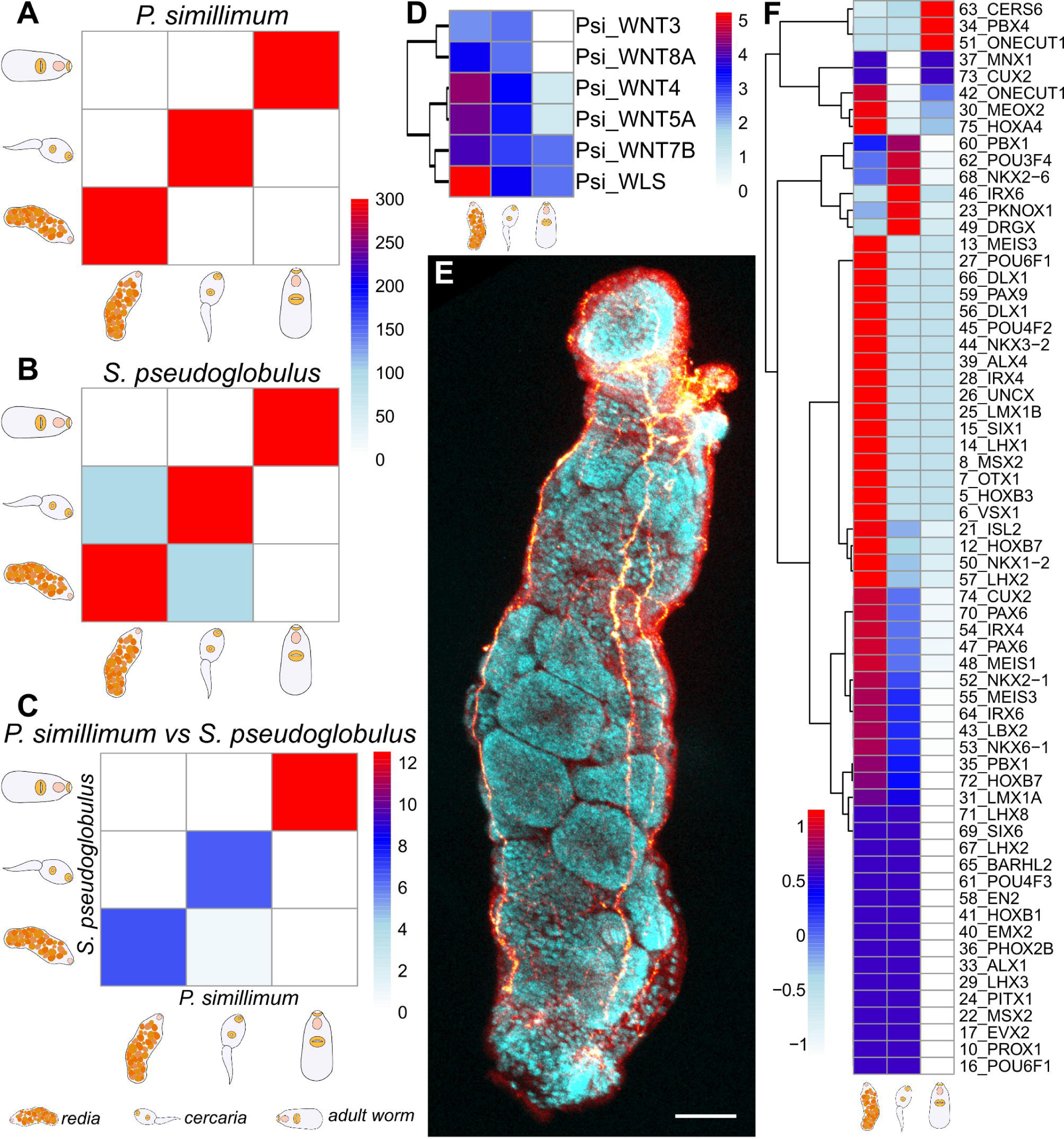
Similarities and differences between molecular signatures of complex life cycles stages. **(A,B)** Overlap measure between *Psilotrema simillimum* **(A)** and *Sphaeridiotrema pseudoglobulus* **(B)** stage-associated gene sets within one complex life cycle. In contrast to *P.simillimum* where stage-associated sets are similar to themselves only **(A)**, in *S.pseudoglobulus* there is a significant overlap between the genes specifically expressed in rediae and cercariae stages **(B)**. **(C)** Overlap measure between *Psilotrema simillimum* and *Sphaeridiotrema pseudoglobulus* stage-associated gene sets. The two-fold differences between Transcriptomes Overlap Measure (TROM) scores obtained for *P.simillimum* and *S.pseudoglobulus* adult worms and for the rediae and cercariae stages of both species is present. A weak “overlap” is also present between *S.pseudoglobulus* rediae and *P.simillimum* cercariae. **(D)** Expression of Wnt family genes in *P.simillimim* life stages. **(E)** Confocal microscopy of *P.simillimum* rediae with developing cercariae embryos. **(F)** Expression of homeobox-containing genes in complex life cycle stages.

Next, we compared how similar are the transcriptomic signatures of the life stages between *Psilotrema* and *Spaeridiotrema*. As a measure of similarity we used expression levels of orthologous genes in each of the stages (TROM score^24^). Our analysis revealed that the transcriptomes of adult worm demonstrate the greatest similarity (TROM-score = 12.53), whereas for cercariae and rediae, the similarity score is almost two times lower and equal to 6.49 and 6.72, respectively. A weak “overlap” is also present between *S.pseudoglobulus* rediae and *P.simillimum* cercariae (TROM-score = 0.82). In total 681 pairs of orthologous genes demonstrate similar expression dynamics in the course of the life cycle progression: 118 pairs have similar expression in the cercariae, 140 in the rediae, and 423 in the adult worms.

In order to obtain a global view of metabolic and signaling pathways that are most important in each of the stages we mapped the sets of stage-specific genes onto KEGG pathways of human (Tables S5-S11). The most “enriched” pathways in *P.simillimum* rediae are those regulating pluripotency of stem cells, Lysosome function, Wnt signaling (see Fig.2D), Axon guidance, Neuroactive ligand-receptor interactions, Adherens junctions, Hippo signaling, Calcium signaling, Cholinergic synapse and Focal adhesion.

Cercaria-specific genes in *P.simillimum* are mainly the members of Metabolic pathways, Calcium signaling, Neuroactive ligand-receptor interaction, cAMP signaling, Lysosome function, Oxytocin signaling, Renin secretion, cGMP-PKG signaling, Adrenergic signaling in cardiomyocytes and Vascular smooth muscle contraction. Last two pathways are rather expectable because a cercaria is the only stage with striated muscles an active locomotion.

The most enriched pathways in *P. simillimum* adult worm are Metabolic pathways, Lysosome, Gap junction, Oocyte meiosis, Galactose metabolism, cGMP-PKG signaling pathway, Oxytocin signaling pathway, Vascular smooth muscle contraction, Apoptosis and Gastric acid secretion.

In general, differentially expressed genes in *Psilotrema* and *Sphaeridiotrema* belong to the similar pathways which is not surprising taking into consideration close phylogenetic relations of these two species (Tables S5-9, 11). However, sets of enriched pathways in both species are overlapping not completely: 32.2% are common in adult worms, 48.6% - in rediae and 65.6% - in cercariae.

One interesting observation is the staggered expression of the homeobox-containing transcription factors during the life cycle of *Psilotrema* and *Sphaeridiotrema* (Fig.2F). It is obvious that some of the homeobox genes are expressed in redia-, cercaria- or adult worm-specific manner. It is remarkable that the majority of homeobox genes are more active in parthenogenetic generations and their expression is down-regulated in the adult worms. This trend might be explained by active morphogenetic processes that take part in during the development of the redia and cercaria. Alternatively, this might indicate that in Trematoda some homeobox genes are utilized as molecular switches regulating transition between different life cycle stages.

## Discussion

Both species selected for our study are characterized by the life cycles with two hosts and the presence of a rediae stage. Due to their phylogenetic positions and well retained ancestral traits these species can be regarded as good models to study the molecular basis the life cycle regulation in the Trematoda. Successful maintenance of *P.simillimum* and *S.pseudoglobulus* under laboratory conditions allowed us to obtain sufficient amounts of material for transcriptomic analysis of all individual stages of their life cycles except for miracidia and metacercaria.

In the absence of genomic references for *P.simillimum* and *S.pseudoglobulus*, their transcriptomes were assembled *de novo* and quality controls suggest that both assemblies are of high quality and completeness (Table 2). The absence of similar sets of the metazoa-specific single-copy orthologs (BUSCOs) in our species and NCBI datasets strongly of other flatworms suggests the evolutionary trend towards considerable gene losses in Trematoda. Probably, observed gene loss is the direct consequence of a parasitic lifestyle. Another explanation for the similar set of “missing” orthologs between our transcriptomes and genomic data may be caused by the absence of their expression in the analyzed stages of the life cycle. In that case, the results of the genome analyses may be compromised by inaccurate *ab initio* identification of coding sequences by gene predicting software. This scenario is corroborated by the observation that on average approximately 23.3% of the BUSCO orthologs from the metazoa_odb9 are “missing” in the analyzed genomes, while only 10.825% of BUSCOs on average are not found in the transcriptomes (Table S1).

**Table 1:** Libraries preparation.

|  | *Psilotrema simillimum* |  |  |  |  |  |
| --- | --- | --- | --- | --- | --- | --- |
| Metrics\Samples | Rediae (1) | Rediae (2) | Cercaria (1) | Cercaria (2) | Adult worm (1) | Adult worm (2) |
| Number of raw reads pairs | 29876923 | 30629840 | 30613174 | 24004011 | 23397182 | 23760802 |
| Human contamination (%) | 0.91% | 0.88% | 0.93% | 1.47% | 0.65% | 0.71% |
| Mollusk contamination (%) | 4.43% | 5.40% | X | X | X | X |
| Chicken contamination (%) | X | X | X | X | 0.99% | 0.29% |
| Trimmomatic survived (%) | 85.96% | 86.31% | 71.59% | 92.74% | 90.91% | 92.90% |
|  | *Sphaeridiotrema pseudoglobulus* |  |  |  |  |  |
| Metrics\Samples | Rediae (1) | Rediae (2) | Cercaria (1) | Cercaria (2) | Adult worm (1) | Adult worm (2) |
| Number of raw reads pairs | 28192806 | 24510459 | 32082446 | 24737749 | 29331490 | 25717702 |
| Human contamination (%) | 0.49% | 0.51% | 0.85% | 0.85% | 0.61% | 0.41% |
| Mollusk contamination (%) | 4.36% | 4.28% | X | X | X | X |
| Chicken contamination (%) | X | X | X | X | 0.70% | 0.22% |
| Trimmomatic survived (%) | 75.94% | 88.93% | 70.45% | 92.79% | 80.22% | 93.16% |

(1|2) – first and second biological replicates, respectively; Human|Mollusk|Chicken – the proportion of potential Human|Mollusk|Chicken-derived read pairs in sample; Trimmomatic survived – the proportion of read pairs remaining after trimming step

**Table 2:** General Statistics of reference transcriptomes quality and completeness measure.

| Metrics \ Assemblies | *P.simillimum* | *S.pseudoglobulus* |
| --- | --- | --- |
| Number of sequences | 163266 | 203659 |
| Number of sequences over 1 kilobase | 28900 | 38618 |
| ESTScan: Number of ORF | 21075 | 49626 |
| N50 (nt) | 917 | 958 |
| BUSCO: Complete and Single (%) | 62.7 | 6.5 |
| BUSCO: Complete and Duplicated (%) | 24.4 | 81.6 |
| BUSCO: Fragmented (%) | 5.1 | 4.0 |
| BUSCO: Missing (%) | 9.0 | 9.3 |
| Fragments mapped (%) | 92 | 89 |
| Good mapping (%) | 81 | 75 |
| Bases uncovered (%) | 4 | 4 |
| Contigs uncovbase (%) | 49 | 50 |
| Contigs uncovered (%) | 5 | 4 |
| Contigs lowcovered (%) | 64 | 50 |
| Contigs segmented (%) | 10 | 13 |
| TransRate assembly score | 0.237 | 0.2095 |
| TransRate optimal score | 0.4566 | 0.4183 |
| Good contigs (%) | 72 | 72 |

BUSCO: Complete and Single | Complete and Duplicated | Fragmented | Missing – the proportion of complete and single-copy | complete and duplicated | fragmented | missing orthologues from Metazoa-odb9 in transcriptomes; Fragments mapped – the total number of read pairs mapping; good mapping – the number of read pairs mapping in a way indicative of good assembly (both members of the pair are aligned; in the correct orientation; on the same contig; without overlapping either end of the contig); Bases uncovered – the propotion of bases that are not covered by any reads; Contigs uncovbase – the proportion of contigs that contain at least one base with no read coverage; Contigs uncovered – the proportion of contigs that have a mean per-base read coverage of < 1; Contigs lowcovered – the proportion of contigs that have a mean per-base read coverage of < 10; Contigs segmented – the proportion of contigs that have >= 50% estimated chance of being segmented; TransRate Assembly score – the geometric mean of all contig scores multiplied by the proportion of input reads that provide positive support for the assembly; TransRate optimal score – TransRate score of the subset of well-assembled contigs; Good contigs – the proportion of contigs with high TransRate scores in transcriptomes.

The results of the comparative gene expression analysis within the life cycles indicate that the majority of genes in trematodes are characterized by a stage-specific expression: 66% in *P.simillimum* and 88% in *S.pseudoglobulus* belong to this category. The absence of significant overlap between the sets of genes with stage-specific expression is clearly noticeable when analyzing *P.simillimum* data (fig.2A). Thus, our approach allows the identification of distinct molecular markers for future analysis of evolutionary trends in each of the stages. In our opinion, the similarities between the gene sets associated with the rediae and cercariae in *S.pseudoglobulus* (fig.2B) are caused by the presence of developing embryos of the cercaria stage inside of the rediae. It is almost impossible to separate them from each other (see Fig.2E). Detected rediae-specific expression of *Wnt* and homeobox-containing genes (fig.2D, F) and results of enrichment analysis (Table S10) may also suggest the active cercariae embryogenesis inside the specimens of the parthenogenetic generation.

Comparison of the expression dynamics throughout the life cycles of two species indicate clear similarities (Fig.2C) but their degree is lower than would be expected from the close phylogenetic relationships of *Psilotrema* and *Spaeridiotrema* within Psilostomatidae. Only 681 pairs of orthologous genes from the total of 7593 OGs found *P.simillimum* and *S.pseudoglobulus* show identical patterns of stage-specific expression indicating that after the separation from the common ancestor a lot of genes have changed their expression patterns.

Two-fold differences between TROM-scores between the adult worms and parthenogenetic stages have several possible implications. Most probably in the course of evolution the expression machinery and gene sets, necessary for amphimictic adult worm formation have been subjected to less changes. As a result, in terms of their transcriptomic signatures, the adult worm of *Psilotrema* and *Sphaeridiotrema* remained much more similar than their corresponding rediae and cercaria stages. What might be the reason for that? It is possible that their definitive hosts (birds) are much more similar than their intermediate hosts (mollusks) and it took much less changes in order to adjust the physiology and morphology of the worms to their hosts.

In both *P.simillimum* and *S.pseudoglobulus*, the majority of genes are transcriptionally active in the adult worms. In *P.simillimum*, the number of sequences associated with this stage differs less than twice from those related with rediae or cercariae. At the same time in *S.pseudoglobulus*, this difference is larger more than 5.5 times. Such striking differences between the adult worms and other stages of *S. pseudoglobulus* life cycle can reflect the diversity and intensity of the processes taking place in the adult worm body as well as may be a result of the presence of several transcriptional activity “sources” in the collected material. We can suggest that the miracidium eggs that are formed in the body of an adult worm as well can be regarded as candidates for these “sources”. Another explanation of this phenomenon may be the presence of cryptic species. Sympatric populations of two morphologically similar species - *S. globulus* and *S. pseudoglobulus*, have already been reported earlier in the study of McLaughlin ^25^ and Bergmame et al. ^26^. The presence of two cryptic species in the *S. pseudoglobulus* population would be a good explanation of the fact that the majority of the orthologues sequences (81.6%, Table 2) from the “metazoan-odb9” database are reported as duplicated in this transcriptome.

Complex life cycle of the trematodes is a system of adaptations ^5^. Every stage has distinct functional contribution to the implementation of the life cycle. Results of the transcriptomic analysis clearly demonstrate the concept of functional complementation among different stages. The major role of the parasitic stage is the increase in the number of parasite individuals at the expense of the host resources. Accordingly, the rediae stage is associated with the pathways involved in protein and RNA synthesis, cell proliferation and cell differentiation (see Wnt signaling pathway, Fig.2D), size regulation during organogenesis (Hippo signaling pathway) and the control of the nervous system formation (Axon guidance). These pathways are also important for the processes associated with cercariae embryogenesis. In the mature worms active assimilation of host resources (Endocytosis; Phagosome; Gastric acid secretion; Ubiquitin mediated proteolysis) and preparation for reproduction (Cell cycle; Oocyte meiosis) are the most important processes. Free-living larvae need to locate the suitable place for encystment. The increased activity of the pathways associated with signal transduction (Neuroactive ligand-receptor interaction; Calcium signaling pathway, cAMP signaling pathway; cGMP-PKG signaling pathway) and muscle contraction (Adrenergic signaling in cardiomyocytes; Vascular smooth muscle contraction), at the cercaria stage in both species could be directly related with the active locomotion and environmental scan.

## Conclusion

Established culturing methods and the availability of transcriptomes provide the basis for future comparative and experimental research and make *P.simillimum* and *S.pseudoglobulus* perspective models to study the evolution of trematode life cycles. While from the current dataset the adult worm stage seems to be the most conserved in term of gene expression profile between two species, comparative data of the miracidium and metacercaria stages as well as qualitative and quantitative differences in the cellular composition of life cycle stages would be important in the future.

## Material and Methods

### RNA isolation and sequencing

Cercariae and parasitic stages (rediae and adult worms), recovered from the hosts, were fixed and stored in the IntactRNA (Eurogene, Moscow, Russia) according to the manufacturer instructions. The stage-specific samples contained approximately 7 adult worms, 100 rediae or 300 cercariae respectively. Prior to RNA isolation, the intactRNA-fixed samples were rinsed in 0.1M phosphate-buffered saline (PBS). The total RNA isolation was done using Quick-RNA™ Microprep kit (R1050, Zymo Research, Irvine, California, USA). For every life cycle stage, two biological replicates were made. The libraries were synthesized using NEBNext Ultra Directional RNA Library Prep Kit for Illumina (E7760, New England BioLabs, Ipswich, Massachusetts, USA). Paired-end sequencing was carried out using Illumina HiSeq 2500 instrument (Illumina, San-Diego, California, USA).

### Libraries analysis

The quality of paired-end read data was manually assessed using FastQC (v0.11.5). Sequencing error correction was performed by Karect (v1.0) (--celltype=diploid –matchtype=hamming) ^27^. All libraries were checked for human tissue contamination (Encode: GRCh38, primary assembly) using BBTools packages (v37.02). Additionally, the reference transcriptome of *Bithynia siamensis goniomphalos* ^28^ and *Gallus gallus* (Encode: *Gallus gallus* v5.0) genome were used for verification of rediae and adult worms libraries host tissue contamination. Sequencing adaptors and nucleotides with a Phred quality score below of 20 removing was performed by Trimmomatic (v0.36) ^29^.

### De novo assembly

For each species the prepared RNA-seq data were pooled together and used to *de novo* assembly of the reference transcriptomes using Trinity (v2.3.2) ^30^. From both transcriptomes only the contigs longer than 200 nucleotides were classified as “good” by the TransRate software (v1.0.1) ^31^ and were selected for farther analysis. Isoform clustering was carried out with CDHIT-EST (-c 0.95) (v4.6) ^32^. The presence of the Metazoan single-copy orthologues in both reference transcriptomes was verified using BUSCO (v3.0.1) (metazoan-odb9, evalue = 1e-3, sp = schistosoma) ^3334^.

The sequences from the latest database may be missing in transcriptomes, both due to the lack of their expression on the analyzed stages and the Platyhelminthes parasitic way of life. Taking this into account, the Metazoan single-copy orthologues searching was also carried out using BUSCO pipeline with same parameters for publicly available *Trichobilharzia regenti* ^23^ and *Opisthorchis felineus* ^22^ transcriptomes, as well as *Schistosoma mansoni* (version 7) ^35^, *S.haematobium* (SchHae_1.0) ^36^, *S.japonicum (*ASM15177v1)^9^, *Opisthorchis viverrini* (*OpiViv1.0*) ^37^, *Clonorchis sinensis* (C_sinensis-2.0)^38^ and *Fasciola hepatica* (Fasciola_10x_pilon)^39^ genomes.

### Aminoacid sequences annotation and orthologues groups searching

The nucleotide sequences were compared with NCBI non-redundant protein database using Diamond BLASTx (v0.9.22; evalue = 1e-5)^40^. Only contigs, that had no match with database or had similarity with sequences from Digenea, were included in further analysis. Their aminoacid sequences identified by the ESTscan ^41^ were re-compared to the NR database using Diamond BLASTp (v0.9.22; evalue = 1e-5).

PANNZER2 (Protein ANNotation with Z-scoRE) service ^42^ was used for the function description and Gene Ontology classes prediction. Only hits of the first rank were taken for further analysis. The functional annotation of the predicted amino acid sequences was carried out with KOBAS3.0^4344^. *Homo sapiens* and *Schistosoma mansoni* were chosen as references.

Reference proteomes of *Schistosoma mansoni* (UP000008854), *Trichobilharzia regenti* (UP000050795), *Opisthorchis viverrini* (UP000054324), *Clonorchis sinensis* (UP00008909), *Fasciola hepatica* (UP000230066) from UniProt database together with predicted aminoacid sequences of *Opisthorchis felineus* ^22^, *Psilotrema simillimum* and *Sphaeridiotrema pseudoglobulus* transcriptomes as well as *Schmidtea mediterranea* (Rhabditophora: Dugesiida) proteins (SmedGD; genome annotations ver.4.0)^45^ were used as input for the OrthoFinder (v2.2.6)^46^.

### Expression level quantification

Salmon (v0.8.2) ^47^ was used for expression level quantification (-l A –gcBias). The analysis was performed on the average of two biological replicates TPM (Transcripts Per Million) values, normalized using the “network centrality analysis” method ^48^. The genes with expression level below 1 TPM in all analyzed stages were excluded from subsequent analysis.

### Life cycle stages comparison

The Jongeneel`s specificity measure ^48^ was used to determine stage-specific genes. Only the genes with positive value (i.e. the gene activity was observed in one stage at a level higher than the sum of all other stages combined) were considered as stage-specific.

The Transcriptome Overlap Measure (TROM) analysis (v1.3) ^24^ was performed to find the similarity between stage-associated genes sets within life cycles and between two species. The stage-associated genes set include stage-specific sequences according to the Jongeneel`s criterion as well as that are active on analyzed, but not all life cycle stages.

The enrichment analysis of the reconstructed pathways with stage-specific genes was carried out using the KOBAS resource (v3.0). The *Homo sapiens* pathways were chosen as the background, and the Fisher`s exact test with Benjamini and Hochberg FDR correction method was used for the statistical analysis. We considered the pathway as “enriched” only when its corrected p-value was below 0.05.

### Immunolabeling and confocal scanning microscopy

For immunostaining studies, the samples were fixed in a paraformaldehyde 4% solution in 0,1 M phosphate buffered saline (PBS) for 8 hours at +4°C, then washed in 0,1 M PBS, incubated in 5% Triton X100 solution in PBS for 24 hours, and blocked in 1% solution of bovine serum albumin in PBS for 6 hours. The blocked specimens were incubated in mixture of rabbit anti-5-HT (S5545, Sigma-Aldrich, St. Louis, USA, diluted 1:1000) and rabbit anti-FMRFamide antibodies (AB15348, EMD Millipore, Burlington, Massachusetts, USA, diluted 1:1000). Rediae were incubated in primary antibody for 24 hours, then washed in PBS with 0,1% Triton X100 and incubated in the secondary anti-rabbit CF488 antibody (SAB4600044, Sigma-Aldrich, diluted 1:1000) for 8 hours. After antibody incubations the samples were washed in PBS and mounted in glycerol. The specimens were examined under Leica TCS SP5 MP confocal laser scanning microscope (Leica Microsystems, Wetzlar, Germany) and analyzed using Fiji software^49^.

### Scanning electron microscopy

For scanning electron microscopy, the animals were fixed in a 2,5% glutaraldehyde solution in 0,05 M sodium cacodilate buffer with a postfixation in 2% osmium tetroxide in 0,05 M sodium cacodilate buffer. The fixed samples were dehydrated in ethanol series of increasing concentration, transferred to anhydrous acetone, critical point dried, sputtered with platinum, and examined using Tescan MIRA3 LMU scanning electron microscope (Tescan, Brno, Czech Republic).

### Availability of data

BioProject has been deposited at NCBI under accession PRJNA516017. *Sphaeridiotrema pseudoglobulus* and *Psilotrema simillimum* Transcriptome Shotgun Assembly projects have been deposited at DDBJ/EMBL/GenBank under the accession GHGK00000000 and GHGL00000000, respectively. The versions described in this paper are the first versions, GHGK01000000 and GHGL01000000, respectively.

## Ethical approval

All applicable international, national, and institutional guidelines for the care and use of animals were followed. We neither used endangered species nor were the investigated animals collected in protected areas.

## Consent for publication

Not applicable

## Authors’ contributions

MAN, VVS, SAD and SVS collected the experimental material. MAN and VVS made the RNA isolations. ARM prepared libraries for RNA-seq and performed the run of the instrument. SAD and SVS carried out the confocal and electron scanning microscopy. MAN and KVK conducted the data analysis. MAN, VVS and KVK wrote the manuscript draft, AIG and AAD contributed substantially to the interpretation of data and to the writing of the manuscript. All authors read and approved the final manuscript.

## Acknowledgements

The scientific research was performed at the Center “Biobank”, Center for molecular and cell technologies, and Center for Culturing Collection of Microorganisms of St. Petersburg State University. Project number in Center for molecular and cell technologies is № 109-9140. The bioinformatics data analysis was performed in part on the equipment of the Bioinformatics Shared Access Center of the Institute of Cytology and Genetics of the Siberian Branch of the Russian Academy of Sciences and using computational resources provided by Resource Center “Computer Center of SPbU” (http://www.cc.spbu.ru/en). MAN wish to thank his parents and close friends for their support and encouragement throughout his study.

## Supplementary materials

Table S1. BUSCO Metazoa_odb9 results

Table S2. BUSCO Common missing

Table S3. OrthoFinder_Statistics

Table S4. OrthoFinder_StatisticPerSpecies

Table S5. Psilotrema_simillimum_rediae_KEGG_enrichment

Table S6. Psilotrema_simillimum_cercariae_KEGG_enrichment

Table S7. Psilotrema_simillimum_adult_worm_KEGG_enrichment

Table S8. Sphaeridiotrema_pseudoglobulus_rediae_KEGG_enrichment

Table S9. Sphaeridiotrema_pseudoglobulus_cercariae_KEGG_enrichment

Table S10. Sphaeridiotrema_pseudoglobulus_rediae_and_cercariae_overlap_KEGG_enrichment

Table S11. Sphaeridiotrema_pseudoglobulsu_adult_worm_KEGG_enrichment

Table S12. Psilotrema_simillimum_combined_results

Table S13. Sphaeridiotrema_pseudoglobulus_combined_results

